# Deciphering binding site conformational variability of substrate promiscuous and specialist enzymes

**DOI:** 10.1101/2024.07.15.603504

**Authors:** Deeksha Thakur, Paras Verma, Shashi Bhushan Pandit

## Abstract

Substrate promiscuity is the ability of enzymes to catalyze reactions with alternate substrate(s) beyond their physiologically relevant ones. Such promiscuous activities expand enzymes functional landscape, enabling evolution or designing novel biochemical reactions. The molecular basis of substrate promiscuity remains elusive, though previous studies have suggested roles of active site structural features, such as flexibility, hydrophobicity, sub-sites, and electrostatics. Moreover, a recent hypothesis proposes the role of active site conformational variability in promiscuity. Accordingly, promiscuous enzymes could accommodate alternate substrate(s) through their pre-existing conformations, whereas specialists have a dominant conformation for their native substrates. To explore the role of active site conformational flexibility in substrate promiscuity, we compared the conformational states of two substrate promiscuous and specialist enzymes by analyzing their binding site structural dynamics from long-time explicit solvent molecular dynamics. Using tICA, we generated conformational states from simulations of the holo-conformation of enzymes. In addition to visual analysis of the variability of binding site conformation states, we performed quantitative estimation of the same using the native functionality score, which measures the contact similarity of conformations to the native structure. We observed that both specialist and generalist enzymes exhibited varied numbers of substrate binding site competent states, indicating that conformational flexibility to accept alternate substrates could exist in both groups of enzymes. Further, it caters to the view of substrate promiscuity as a continuum feature of enzymes.

## Introduction

Enzymes are biocatalysts known for their remarkable specificity towards respective cognate substrate(s). However, some enzymes exhibit promiscuous behavior, being able to catalyze alternate substrates (substrate promiscuity) or catalyze different reactions (catalytic promiscuity) utilizing the same active site (Copley, 2017; Khersonsky & Tawfik, 2010). Although typically characterized by their low catalytic efficiency, these undesirable secondary promiscuous activities serve as reservoirs for evolving novel biochemical transformations, which can be essential for an organism’s metabolism in response to genetic or environmental perturbations (D’Ari & Casadesús, 1998; Guzman et al., 2019). Indeed, promiscuous enzymes are widely distributed in organisms, catalyzing various reactions (Martinez-Nunez, Rodriguez-Escamilla, Rodriguez-Vazquez, & Perez-Rueda, 2017; Nam et al., 2012). Besides the physiological significance of enzyme promiscuity, it has broader applications in developing novel biocatalysts exploiting catalytic promiscuity features and metabolizing non-native substrates to degrade harmful chemical compounds (Aharoni et al., 2005; Kovacs, Szappanos, Tengolics, Notebaart, & Papp, 2022). This ability to intentionally and efficiently expand the substrate range of promiscuous enzymes (generalist) has thus become a focal point in enzyme engineering (Agarwal, Bernard, Bafna, & Doucet, 2020; Copley, 2017; Dhamankar & Prather, 2011).

Understanding the molecular basis of substrate promiscuity can facilitate the rational design of enzymes for catalyzing desired substrates (Garrabou, Beck, & Hilvert, 2015). Previous studies have highlighted the roles of active site flexibility, hydrophobicity, subsites, electrostatics, and metal ions in accommodating alternate substrates for catalysis among promiscuous enzymes (Babtie, Tokuriki, & Hollfelder, 2010; Copley, 2012, 2017; Khersonsky & Tawfik, 2010). However, our recent work has found that active site structural features are similar between promiscuous and specialist enzymes (substrate-specific), suggesting a continuum of substrate promiscuity having two extremes ranging from multi-specific to highly substrate-specific enzymes (Thakur & Pandit, 2022). Most of these characteristic features of promiscuous/specialist enzymes are primarily uncovered or understood from a static state of an enzyme conformation captured in the crystal structure. However, enzymes undergo a range of motions, from local active site fluctuations and loop motions to global conformational changes required for substrate binding, chemical catalysis, and product dissociation (Corbella, Pinto, & Kamerlin, 2023; Lemay-St-Denis, Doucet, & Pelletier, 2022; Maria-Solano, Serrano-Hervas, Romero-Rivera, Iglesias-Fernandez, & Osuna, 2018; Singh, Fenwick, Dyson, & Wright, 2021; Wrabl et al., 2011). Many recent studies have demonstrated the roles of protein dynamics in catalysis, catalytic efficiency optimization, and the emergence of new catalytic functions through enriching alternate conformational sub-states (E. C. Campbell et al., 2018; Fraser et al., 2009; Romero-Rivera, Garcia-Borras, & Osuna, 2017). The extensive studies on β-lactamase ancestral enzyme reconstruction (Kolbaba-Kartchner, Kazan, Mills, & Ozkan, 2021; Risso et al., 2017) and directed evolution of enzymes (E. C. Campbell et al., 2018; Tokuriki et al., 2012) highlight the potential role of conformational dynamics in molecular evolution (Hart, Ho, Dutta, Gross, & Bowman, 2016; Osuna, Jimenez-Oses, Noey, & Houk, 2015; Petrovic, Risso, Kamerlin, & Sanchez-Ruiz, 2018), suggesting that equilibrium among sub-states could be modulated by mutagenesis (E. Campbell et al., 2016; Serrano-Hervas, Casadevall, Garcia-Borras, Feixas, & Osuna, 2018; Wang, Liu, Zhu, Peng, & Ma, 2023). It is significant to mention that evolution of enzymes strikes a balance between conformational flexibility and its function (Aharoni et al., 2005; Tokuriki et al., 2012).

The conformational selection hypothesis proposes the pre-existence of conformation in the ensemble primed to accept a particular substrate (Atkins, 2015, 2020). It has also been speculated that the presence of a substrate shifts the equilibrium towards the conformational cluster that favors its binding (Honaker, Acchione, Zhang, Mannervik, & Atkins, 2013). Considering these features essential for enzyme function, it has been hypothesized that conformational energy landscape of specialist enzymes consists of the energetic minimum state populated for a particular substrate binding and catalysis. In contrast, promiscuous enzymes have several conformational minima accommodating catalysis of alternate substrates while having near-native active site geometry (Copley, 2017; Damry & Jackson, 2021; Khersonsky & Tawfik, 2010; Maria-Solano et al., 2018). Most studies have simulated promiscuous (Romero-Rivera et al., 2017), laboratory (directed) evolution or mutant enzymes to explore conformational space (Behiry et al., 2018; Corbella et al., 2023; Romero-Rivera et al., 2017), and similar studies on specialist enzymes are somewhat limited. We performed long-time molecular dynamics simulations to investigate variations in the structural dynamics between active sites of specialist and generalist enzymes. The simulation trajectories were subjected to time-structure independent component (tICA) analysis followed by clustering to identify representative conformational states. Apart from other measures to compute near-native active site geometries, we devised a native functionality score, which is a step function to compute spatial contacts similarity of active site residues between structures. Here, we present the case study of four enzymes (two from each category) to support our argument of the existence of a broad extent of binding site variability in both the enzyme groups.

## Results

To explore conformational basis of enzyme promiscuity, we investigated the binding site (BS) conformational flexibility of two enzymes each from promiscuous and specialist groups, which we had previously curated, for comparison and contrast (Thakur & Pandit, 2022). Cytidine deaminase and dihydrofolate reductase were chosen in the promiscuous category and erythritol kinase and p-hydroxybenzoate reductase were chosen to represent the specialist category. To analyse the conformational variability, we performed long-time (1.5 μs) equilibrium Molecular Dynamics (MD) simulation of holo-state enzyme structure and analyzed their representative conformational sub-states using various structural similarity measures. As we need to recognize structure closely related to binding/catalytic competent conformation for assessing their ability to accommodate cognate substrate, the representative conformation states from simulation were compared to the BS of native structure. Based on previous studies, we hypothesized that promiscuous enzymes would exhibit multiple conformers in comparison to specialist enzymes. The conformational states obtained as cluster representatives after clustering of trajectories on binding site residues (BSR) interatomic distance as features. Apart from using traditional measure of structural similarity, RMSD (Root Mean Square Deviation), we devised a measure called native functionality score (*fχ*), to assess competent state of substrate binding based on interatomic distances of BSR. To avoid any bias to an atom, equal weightage is associated with all interatomic distances. The *fχ* is a modified chi-value score from the work of Nissley et al., 2022 that was used for measuring native state in folding simulations. The native functionality score (*fχ*) ranges from 0 to 1, with higher score suggestive of similarity to native structure, while lower score defines loss of interatomic contacts (shown to be important in the crystal structure for binding to the native substrate). Below we discuss conformation variations of promiscuous and specialist enzymes.

## Promiscuous enzymes

### Cytidine deaminase (CDA)

Cytidine deaminase enzyme recycles free pyrimidines and is involved in both anabolic (synthesis of nucleotides for incorporating in DNA/RNA) and catabolic (degradation of pyrimidines as a supply of carbon and nitrogen) metabolic processes (Frances & Cordelier, 2020). This enzyme belongs to the cytidine and deoxycytidylate deaminase family, which requires zinc ion for its enzymatic activity. CDA sequence is conserved across organisms from prokaryotes to eukaryotes with *E. coli* CDA (EcCDA) sharing 39% sequence identity with human (hCDA) sequence (Frances & Cordelier, 2020). The tertiary structure of EcCDA contains CMP (cytidine monophosphate)/dCMP (deoxycytidine monophosphate)-type deaminase domain composed of a central β-sheet with one or more α-helices on each side, which is characteristic feature of APOBEC family of enzymes (Reizer, Buskirk, Bairoch, Reizer, & Saier, 1994). Further structural and mechanistic studies of EcCDA, *Klebsiella pneumonia* (KpCDA) and other Zn^2+^ dependent CDA enzymes indicated that the loop motion in the binding site could mediate the catalytic activity and substrate specificity for different substrates (Marx, Galilee, & Alian, 2015; Rathore et al., 2013). EcCDA apart from cytidine (its native substrate) can catalyze 2′-deoxycytidine (Hosono & Kuno, 1973) exhibiting substrate promiscuity.

We investigated the conformational variability of the EcCDA binding site region to understand the local structural dynamics, which could facilitate accommodation of alternate substrates. We performed a long-time (1.5μs) explicit solvent equilibrium MD simulation of EcCDA enzyme structure (PDB: 1cttA). The structure is bound to a transition state analogue (3,4-Dihydro-1H-pyrimidin-2-one nucleoside), which defined the substrate binding and Zn ion binding sites (Xiang, Short, Wolfenden, & Carter, 1995). As discussed in the methods, holo-enzyme structure without any ligand/metal is used for MD simulations to extensively explore the conformational space close to substrate-analogue bound state. The simulation trajectories were analyzed for residue fluctuations using Root Mean Square Fluctuation (RMSF) followed by identifying various dominant conformational states using approaches described in methods section. Overall structure of EcCDA enzyme remained stable during simulation (Figure S1A). Subsequently, we analyzed backbone (Cα) residue fluctuations, using RMSF as a measure shown in Figure 1A, which also highlights the binding/catalytic residues. As is evident, the highest fluctuations are primarily observed in the loop regions (loop^61-73^, loop^93-99^ and loop^159-174^). Of these, residues in the loop^93-99^ and loop^159-174^ forms inter-protomer interacting interface, indicating that this interaction probably is essential for stabilizing these loop conformations and absence of it contributes to high mobility. Their structural dynamics is also apparent in smoothened representation of these loops (Figure 1B). The loop^61-73^ forms the binding site and it harbours two conserved residues S69 and F71 showing maximum fluctuation (Figure 1A), which interacts with E91 and N89 respectively (Figure S1B). These residues in turn are hydrogen bonded to O3′ atom of the substrate analogue (DHZ). Moreover, a separate study on KpCDA enzyme structure has found that S69 and F71 interacting with bound water molecule (Liu et al., 2019). Thus, indicating that loop^61-73^ residues play an important role in binding and probably in catalysis as well (Liu et al., 2019). Other binding residues, unlike these, exhibit limited mobility except residues C129 and C132, which coordinates with Zn ion.

**Figure 1:**
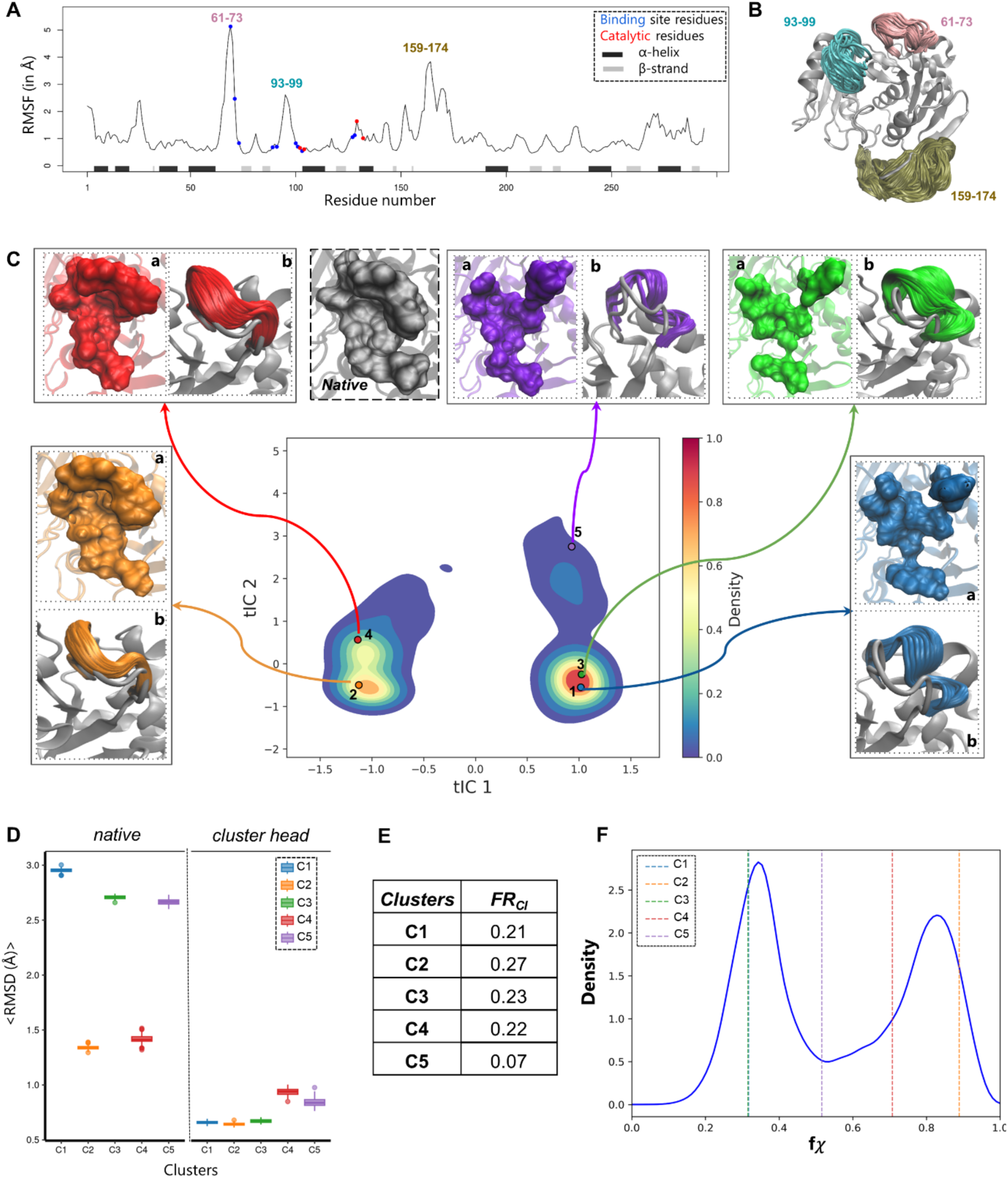
Analysis of cytidine deaminase (PDB:1cttA) MD simulations: (**A)** Line plot showing RMSF values of Cα atoms from MD simulation trajectory (30,000 frames). Binding site and catalytic residues are shown on the plot in blue and red color respectively. (**B)** Tertiary structure of enzyme with highly mobile loops shown as smoothened representation (see methods). (**C)** tICA plot showing projection of MD trajectories as conformational density on first two tIC components. The five cluster representatives (numbered from 1 to 5) after MiniBatchKmediods clustering is projected on the tICA plot with each of them colored differently. For each cluster, the binding pocket is illustrated with surface representation in sub-panel (a) and smoothened representation of highly variable region of the loop is shown in sub-panel (b). The native active site in surface representation is shown in grey. (**D)** The left panel (*native*) shows box plot of mean RMSD of sampled conformations to the first frame of simulation and right panel (*cluster head*) is box plot of mean RMSD of sampled conformations to the cluster representative. (**E)** Tabulated fraction of members (FR_cl_) for each cluster with respect to total frames (see methods)**. (F)** The native functionality (*fχ*) density plot is shown in blue curve and the mean of each cluster sampled mean is shown in dotted line.

Having analyzed the local backbone fluctuations, we examined detailed conformational landscape spanned by the binding site residues. For this, we featurized interatomic distances of the closest heavy atoms of BSR for dimensional data reduction method using tICA (see methods) and projected conformations on reduced independent component vectors (tICs). The projection of trajectories on first two tICs show distinct regions (Figure 1C) characterizing broadly two conformational regions. Further, we clustered trajectories based on interatomic distances (BSR) into five clusters to characterize conformational states of EcCDA (Figure 1C), which shows some states lie in close vicinity of each other, suggesting similarity in their binding site region. As we have clustered structures based on BSR, we first assessed their structural variation by superposing BSR of cluster members on the native structure (Figure 1D). We sampled clusters to quantitatively assess BSR features (see methods). For validating the consistency of structural variability of binding site, the cluster members were superposed on cluster representative structure (see methods). It can be seen (Figure 1D), the clusters C2 and C4 are closer to native structures with average ∼1.4 Å RMSD and rest others exhibit relatively poorly tuned binding site. Overall cluster members are consistent in their binding site conformation as exhibited with their average low RMSDs (Figure 1D) to the cluster representative structure. For visual analysis of binding site conformations, we compared cluster representative binding pocket to the native structure (Figure 1C, panel (a) of each cluster) that showed that clusters 2 and 4 are similar to the native binding pocket, as shown in surface representation. However, rest other clusters exhibit relatively open binding site (Figure 1C) due to loop motion indicating that it may not be able to tightly accommodate native substrates. Subsequently, we also examined the loop regions constituting binding site and observed the loop^61-73^ motion segregates into broad two distinct conformations with one being shown in C1, C3 and C5, which are spatially distant from native leading to a more open state (Figure 1C, panels (b) of each cluster). On the contrary, C2 and C4 loops are native-like and retain binding site competent conformation. It is important to note that C2 and C4 together constitute nearly half of the conformation population (Figure 1E). Other conformational states may possibly accommodate larger substrate as the loop is largely in open state. However, it is imperative to note that interactions of N89 and E91 with loop (S69 and F71) (figure S1B) is disrupted in these open states, which may pose challenge in accommodating other larger compound with tight binding.

As spatial orientation of BS atoms is essential for stable substrate interactions, we quantified this using a step function computed as native functionality score (*fχ*), which measures variation of atom positions relative to the reference state (native structure) with an allowed limit (see methods). The C2 and C4 clusters are visually native-like states, which is also evident from variability score as members of these have the highest average *fχ* scores, while rest other members of clusters (C1, C3 and C5) show varying score indicating differential degree of similarity to native binding site atom orientations (Figure 1F). The complete distribution of *fχ* of all the sampled conformations is shown in Figure S1C.

The equilibrium simulation showed that EcCDA exhibit conformational variability suggesting that there are distinct conformational states, mostly due to the fluctuation of loop ^61-73^ indicating its possible role in accommodating alternate substrate(s). The presence of conformations with varied native functionality might contribute to the accommodation of alternate substrates, thus contributing to the enzyme’s ability to catalyze multiple substrates.

### Dihydrofolate reductase (DHFR)

Next, we analyzed binding site variability of another promiscuous enzyme human dihydrofolate reductase (hDHFR), which is a pivotal enzyme in the folate metabolic pathway. It is an essential enzyme in the synthesis of tetrahydrofolate. As tetrahydrofolate is critical for biosynthesis of purines, thymidylate, and certain amino acids, it has been shown that cytosolic hDHFR is upregulated in many types of cancer (Davies et al., 1990)(Zheng et al., 2018). Moreover, this enzyme inhibition in cancer cells leads to disruption of biosynthesis of essential metabolites followed by cell death, thus, making hDHRF attractive drug (antifolates) target in cancer treatment (Zheng et al., 2018). The enzyme tertiary structure consists of central 8-stranded β-pleated sheet with surrounding α-helices, which mostly form the active site.

As has been described before, we analyzed the binding site dynamics of hDHFR from long-time explicit solvent equilibrium MD simulation of holo-conformation of the enzyme (PDB: 1dhfA). Overall, the enzyme remained stable throughout the simulation and the same can be observed in the RMSD plot (figure S2A). Initially, we focused on backbone dynamics measured using RMSF (Figure 2A) and as can be seen most fluctuations are in the loop regions and C-terminal region of the protein. (Figure 2A). The loop regions loop^19-25^, and loop^40-45^ have extensive variations in comparison to loop^61-68^, which also harbors BSR and the same can be visually observed in smoothened representations (Figure 2B). To examine the structural dynamics of BSR, following the procedure discussed before, we used heavy atom interatomic distances as features in tICA followed by clustering of trajectories in five clusters (see methods). The projection of conformations along the first two tICs components revealed a dense central region (figure 2C) connected with minor small, populated states suggesting limited variations of BS regions. The distribution of members in five clusters are shown in Figure 2E having most members belonging to C1 and C2. The cluster representative binding pockets are visually (surface representation) similar to the one in native state (Figure 2C, panel (a) of each cluster) with minor variations occurring due to loop^61-68^ motion that lies to the periphery of the pocket. Importantly, the interior cavity remained largely unchanged, which is essential for binding substrate. The loop^61-68^ variability is mostly uniform across all clusters and does not distinguish among pockets in clusters (loop smoothened panels (b) in Figure 2C). The average RMSD of clusters to native structure is below 1.5 Å, which is expected as variations in BSR are limited (Figure 2D). The consistency of clustering is also evident from average low RMSDs of members to their cluster representative (Figure 2D).

**Figure 2:**
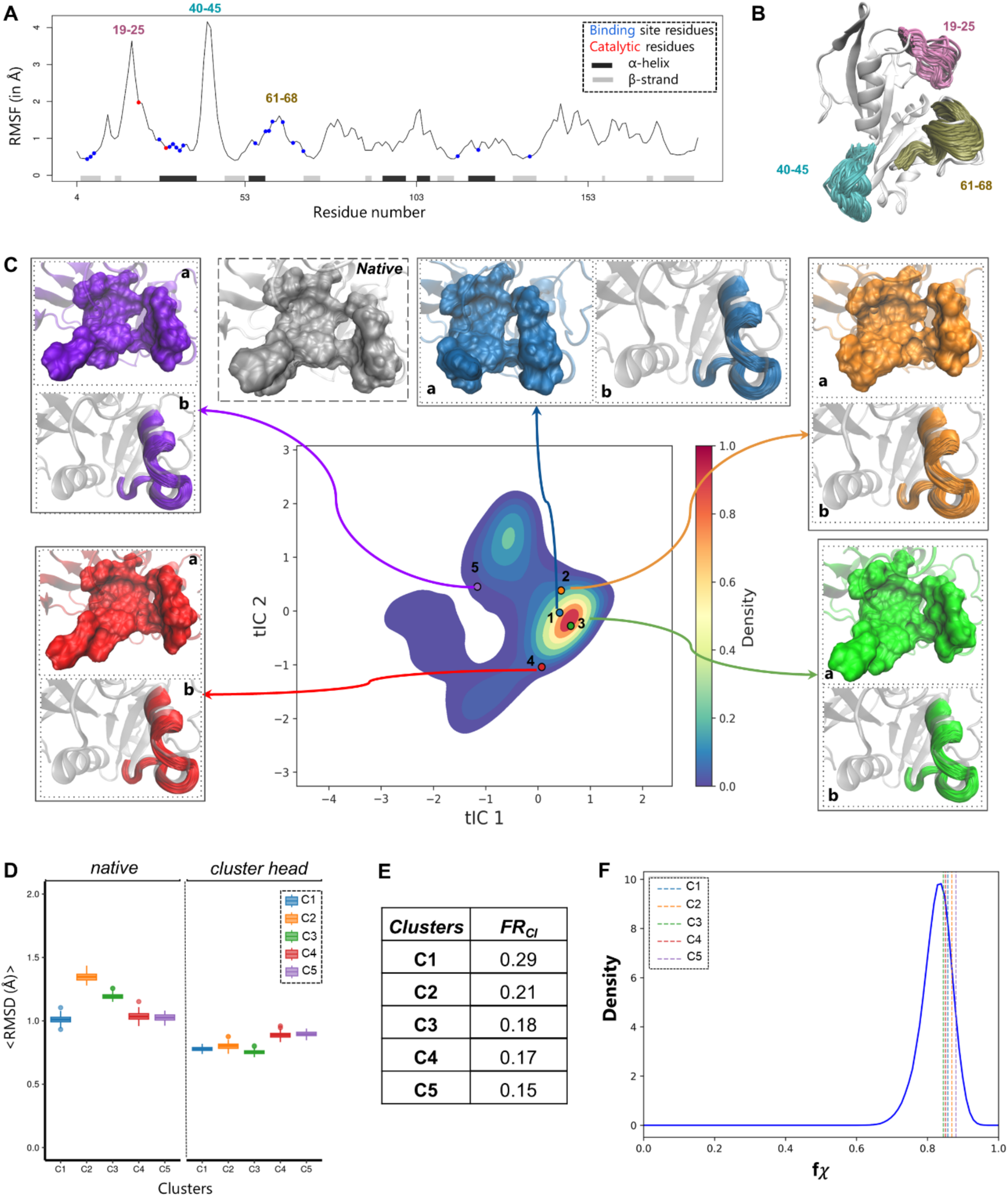
Analysis of dihydrofolate reductase (PDB:1dhfA) MD simulations: (**A)** Line plot showing RMSF values of Cα atoms from MD simulation trajectory (30,000 frames). Binding site and catalytic residues are shown on the plot in blue and red color respectively. (**B)** Tertiary structure of enzyme with highly mobile loops shown as smoothened representation (see methods). (**C)** tICA plot showing projection of MD trajectories as conformational density on first two tIC components. The five cluster representatives (numbered from 1 to 5) after MiniBatchKmediods clustering is projected on the tICA plot with each of them colored differently. For each cluster, the binding pocket is illustrated with surface representation in sub-panel (a) and smoothened representation of highly variable region of the loop is shown in sub-panel (b). The native active site in surface representation is shown in grey. (**D)** The left panel (*native*) shows box plot of mean RMSD of sampled conformations to the first frame of simulation and right panel (*cluster head*) is box plot of mean RMSD of sampled conformations to the cluster representative. (**E)** Tabulated fraction of members (FR_cl_) for each cluster with respect to total frames (see methods)**. (F)** The native functionality (*fχ*) density plot is shown in blue curve and the mean of each cluster sampled mean is shown in dotted line.

Subsequent to structural evaluation, we computed native functionality score (Figure 2F) and consistent with previous observations, the mean *fχ* score of sampled members is above 0.8 and distribution varies from 0.7 – 0.9 (Figure S2B) indicating that interatomic distances are mostly maintained across clusters. The overlapping distribution of average *fχ* scores for all the sampled conformations for each cluster can also be seen in Figure S2B. These conformational variability analysis of hDHFR showed that BS is mostly similar with variations lying primarily in the periphery of the binding pocket (loop^61-68^) suggesting this promiscuous enzyme largely exhibit similar BS conformations unlike as has been seen in another promiscuous enzyme (EcCDA). Moreover, it is likely that small loop variation of loops probably accommodates alternate substrate as it does not disrupt the binding pocket.

## Specialist enzymes

### 4-diphosphocytidyl-2-C-methyl-D-erythritol kinase (CDP-ME Kinase)

We selected *E. coli* 4-diphosphocytidyl-2-C-methyl-D-erythritol kinase from the specialist enzyme group for analyzing BS conformational variability. CPD-ME kinase is an important enzyme in the nonmevalonate biosynthetic pathway of isopentenyl diphosphate and dimethylallyl diphosphate. The CDP-ME kinase subunit consists of ∼40% α-helices, 28% β-strands, and two short 3_10_-helices, which form a two-domain fold, typical of the GHMP kinase superfamily. The cofactor or ATP-binding domain is made up four-stranded β-sheet on one side and a bundle of five α-helices on the other. The substrate-binding domain constitutes residues from two twisted antiparallel β-strands and five α-helices positioned around the periphery. It catalyzes ATP-dependent phosphorylation that results in 4-diphosphocytidyl-2C-methyl-d-erythritol-2-phosphate (Miallau et al., 2003). As this enzyme is absent in mammals but essential in bacteria, it is an attractive drug target for development of broad spectrum anti-microbial drug (Eoh et al., 2009).

We performed long-time MD simulation of CDP-ME kinase (PDB: 1oj4A) to examine binding site conformation variability (see methods). The structure remained mostly stable during the simulations as is seen in the RMSD (Figure S3A) plot with respect to simulation time. Upon analyzing backbone fluctuations using RMSF, we observed three regions undergoing large motions (loop^23-28^, loop^155-158^, and loop^176-187^) apart from N/C-terminal of the protein (Figure 3A). Notably, almost all these highly mobile regions encompass BSR with some of them showing fluctuation more than 3Å. The extent of mobility is also depicted using smoothened representation of trajectories in Figure 3B. The binding site residue on loop^156-158^ (V156) interacts with the residues (H26 and Y25) from another loop^23-28^, which lies in inter-protomer interface region. The absence of inter-protomer stabilizing interaction of the β-turn (21-25) increases its mobility, thereby affecting the interaction of it with the V156 and affecting the loop^155-158^. We further examined BSR dynamics using the procedure as has been described before of considering interatomic distances of heavy atoms of BSR. The conformations projected on the top two tICs showed two regions with the one having large spread (Figure 3C) and consisting of all five cluster centers lying on it. The structural dynamics of clustered structures were analyzed using representative structure and it is evident that binding pocket (surface representation of pocket) is poorly formed across clusters in comparison to the native, although C2 and C3 seems to maintain some feature of the pocket; C2 retained overall shape but the pocket size has decreased in it due to inward motion of loop^23-28^, where C3 has more open binding pocket (Figure 3C, panels (a) of each cluster). Among clusters, C4 has the most disrupted binding, which is also evident from its highest average RMSD to the native site. The loss of binding pocket similarity is also reflected in the higher (>1.5 Å) average RMSD of sampled cluster members to native across clusters (Figure 4D). The cluster members show consistent average low RMSD between cluster representative and its respective members (Figure 4D).

**Figure 3:**
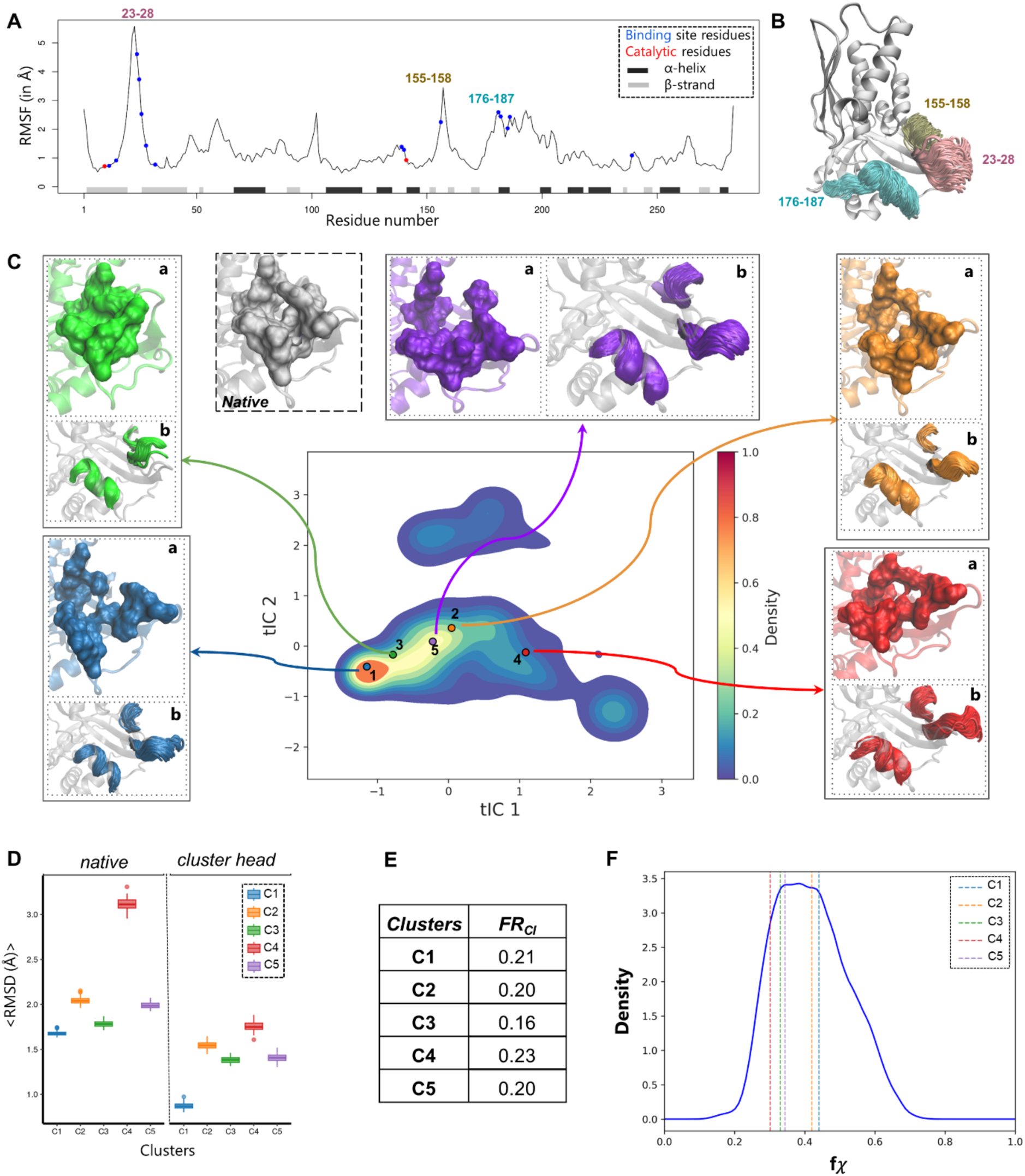
Analysis of 4-diphosphocytidyl-2-C-methyl-D-erythritol kinase (PDB:1oj4A) MD simulations: (**A)** Line plot showing RMSF values of Cα atoms from MD simulation trajectory (30,000 frames). Binding site and catalytic residues are shown on the plot in blue and red color respectively. (**B)** Tertiary structure of enzyme with highly mobile loops shown as smoothened representation (see methods). (**C)** tICA plot showing projection of MD trajectories as conformational density on first two tIC components. The five cluster representatives (numbered from 1 to 5) after MiniBatchKmediods clustering is projected on the tICA plot with each of them colored differently. For each cluster, the binding pocket is illustrated with surface representation in sub-panel (a) and smoothened representation of highly variable region of the loop is shown in sub-panel (b). The native active site in surface representation is shown in grey. (**D)** The left panel (*native*) shows box plot of mean RMSD of sampled conformations to the first frame of simulation and right panel (*cluster head*) is box plot of mean RMSD of sampled conformations to the cluster representative. (**E)** Tabulated fraction of members (FR_cl_) for each cluster with respect to total frames (see methods)**. (F)** The native functionality (*fχ*) density plot is shown in blue curve and the mean of each cluster sampled mean is shown in dotted line.

**Figure 4:**
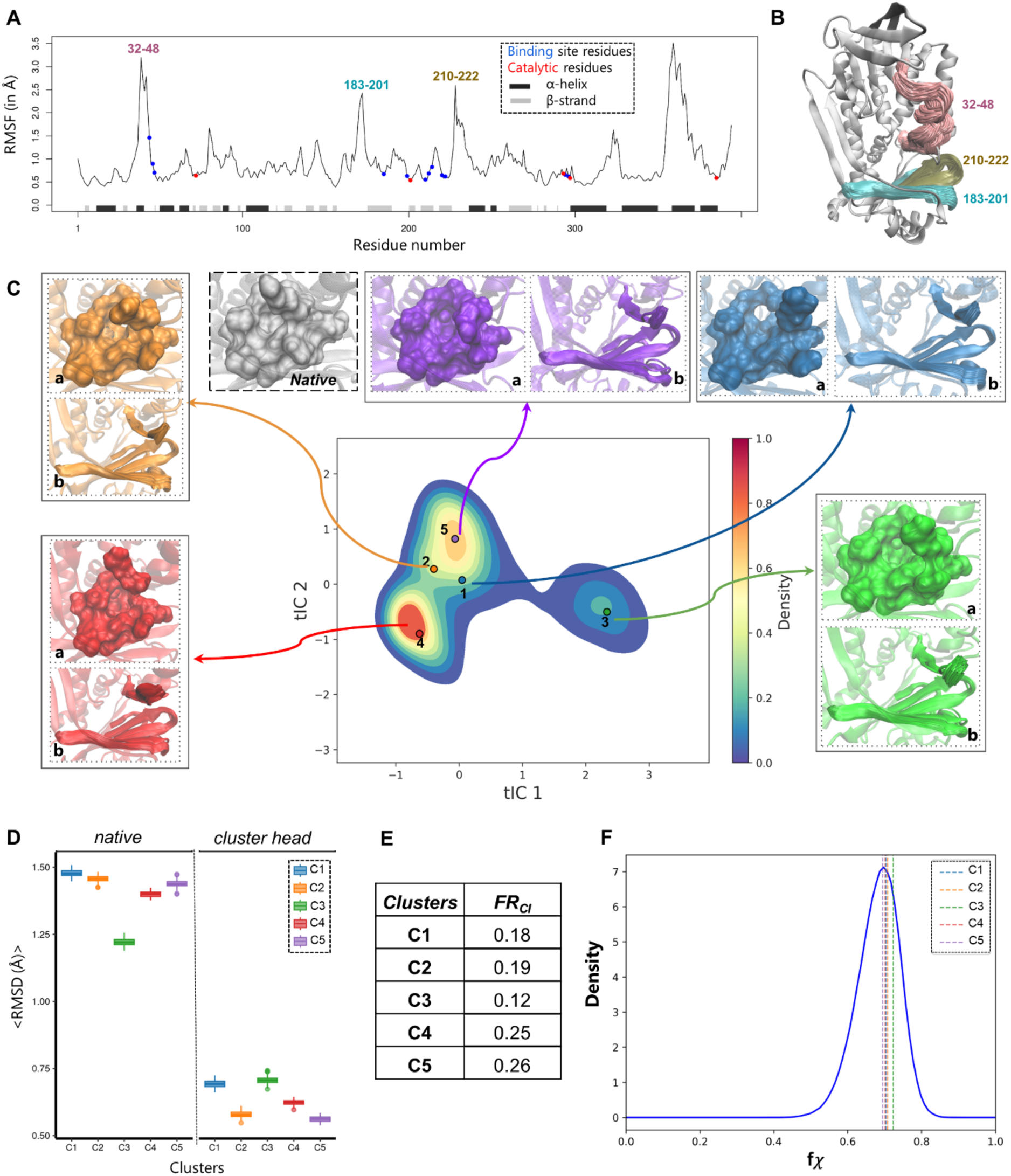
Analysis of p-hydroxybenzoate hydroxylase (PDB:1docA) MD simulations: (**A)** Line plot showing RMSF values of Cα atoms from MD simulation trajectory (30,000 frames). Binding site and catalytic residues are shown on the plot in blue and red color respectively. (**B)** Tertiary structure of enzyme with highly mobile loops shown as smoothened representation (see methods). (**C)** tICA plot showing projection of MD trajectories as conformational density on first two tIC components. The five cluster representatives (numbered from 1 to 5) after MiniBatchKmediods clustering is projected on the tICA plot with each of them colored differently. For each cluster, the binding pocket is illustrated with surface representation in sub-panel (a) and smoothened representation of highly variable region of the loop is shown in sub-panel (b). The native active site in surface representation is shown in grey. (**D)** The left panel (*native*) shows box plot of mean RMSD of sampled conformations to the first frame of simulation and right panel (*cluster head*) is box plot of mean RMSD of sampled conformations to the cluster representative. (**E)** Tabulated fraction of members (FR_cl_) for each cluster with respect to total frames (see methods)**. (F)** The native functionality (*fχ*) density plot is shown in blue curve and the mean of each cluster sampled mean is shown in dotted line.

The quantitative assessment of binding site conformational variability using the average *fχ* score shows overall values <0.5 suggestive of poor structural orientation of BSR in cluster members. This is consistent with the visual interpretation of the conformational variations observed during simulations. The distribution of *fχ* score of sampled conformations is shown in Figure S3B. Upon detailed analysis of the interatomic contacts in BSR revealed that apart from hypermobility of loop regions, flipping of binding site residue F32 (Figure S3C), which interacts with many other atoms, affects the *fχ* score.

CDP-ME kinase despite being a specialist enzyme with substrate specificity substrate, showed large conformational diversity of BSR, of which many of these are not close to native state. The present analysis indicates there is a possibility of more than one conformation, which could accommodate different substrates. It is plausible that enzyme requires bound cofactor for generating near-native conformations as some of the loop region involved in high mobility also interacts with cofactor.

## p-hydroxybenzoate hydroxylase (*PBH*)

We considered *Psuedomonas aeruginosa* para-hydroxybenzoate hydroxylase as the second case study for specialist enzyme. It catalyzes the insertion of oxygen into substrates by means of the labile intermediate, flavin C(4a)-hydroperoxide (Gatti et al., 1994) and catalyzes the incorporation of an atom of dioxygen into p-hydroxybenzoate to form 3,4-dihydroxybenzoate in a two part reaction (Entsch & Ballou, 1989). The tertiary structure of enzyme (PDB: 1docA) is composed of three domains: a large, globular domain (residues 1-180), where the flavin cofactor binds; a smaller domain (residues 180-270) having a single helix and an antiparallel β-sheet; and a third, entirely helical domain (residues 270-394). The active site is located at the interface of three domains (Wang et al., 2002).

We performed long-time MD simulations of PBH enzyme (PDB: 1docA) to explore BSR conformations. The structure remained stable throughout the simulation (Figure S4A). On analyzing the fluctuation using RMSF, active site residues showed small mobility (figure 4A), except for few BSR lying on a mobile loop^32-48^ (figure 4B). This suggested that BSR does not undergo large structural variations. Subsequently, projection of the conformations on reduced dimensions using tICA on the first two tIC components revealed a conformational density spread across two regions (Figure 4C). The structural similarity assessed using RMSD of sampled conformations showed average low RMSD (<1.5 Å) and the cluster C3 closest to the native (1.2 Å) (Figure 4D). The cluster members consistently showed BSR similarity to the cluster representative as shown by average low RMSD (<0.75 Å) (Figure 4D). The comparison of binding site pocket of cluster representatives with native showed similarity between them, with changes mostly to the periphery of the pockets (Figure 4C, panels (a) of each cluster). The fluctuations of the loop regions closed to the BS also have minimal variability as seen in their smoothened representations (Figure 4C, panels (b) of each cluster).

Next, we also compared the native functionality score (*fχ* score) of the sampled conformations, which showed average score >0.7 (Figure 4F) indicating that BSR atomic spatial orientations of clusters are like that of native structure. Overlapping distributions of all the sampled conformations can be seen in figure S4B. One of the BSR, R44 which resides on the aforementioned loop^32-48^, exhibits a large side chain motion which likely affects the native functionality score as all the cluster heads have different side chain orientation as compared to that of native (in grey) and each other (Figure S4C). This residue makes multiple interactions with the ligand and other BSR, using both its side chain and backbone atoms. Except this, rest of the BSR remain spatially constrained across all clusters. This specialist enzyme represents the conformational ensemble having binding site competent states in all clusters.

Through these case studies, we examined both specialist and promiscuous enzymes for their conformational variability of BSR highlighting varying degrees of structural and binding site competent states, as assessed by native functionality score. This shows that the ability of enzymes to accommodate alternate substrates may be prevalent in specialist and promiscuous enzymes alike. Growing research on substrate promiscuity also suggests that it could never be fully realized because of the absence of the knowledge of the conditions in which these latent activities can be observed (Copley, Newton, & Widney, 2023). This argues in favour of substrate promiscuity being a continuum feature and its extent could be an enzyme specific property, linked to its biological role in the cell. Understanding the conditions under which latent activities can be observed is crucial for fully realizing substrate promiscuity, emphasizing that it is a nuanced and enzyme-specific property.

## Conclusion and discussion

In the current work, we examined the conformational basis of enzyme promiscuity and determined whether promiscuous enzymes exhibit a greater conformational flexibility of BSR for accommodating alternate substrates. Towards this, we analyzed the conformational variability of BSR of promiscuous and specialist enzymes through long-time equilibrium MD simulations and examined conformational states using tICA followed by clustering of them. The case studies of two enzymes from each category demonstrated that BSR conformational variations as assessed visually or quantitatively using native functionality score, is observed in both categories of enzymes. In specialist enzyme (CPD-ME Kinase), the diverse binding pockets are generated, and it is difficult to ascertain which of these can accommodate native substrate. Thus, our findings suggest that binding site variability is present in both groups of enzymes.

Building on our previous work, which demonstrated structural similarities between promiscuous and specialist enzymes, we proposed that substrate promiscuity is a continuum feature. The present study reinforces our hypothesis also for conformational basis of promiscuity through four case studies that highlight the native functionality extremes in both enzyme categories. We observed that variability, traditionally attributed to promiscuous enzymes, is also present in specialist enzymes. Conversely, promiscuous enzymes can exhibit low binding site variability while still accepting multiple substrates. We speculate that different enzymes, whether promiscuous or specialist, may exhibit varying degrees of conformational variability influenced by factors such as their cellular roles. This analysis underscores the complexity of enzyme behavior and challenges the conventional distinction between promiscuous and specialist enzymes, highlighting the nuanced continuum of substrate specificity and binding site variability.

## Methods

To study conformational diversity of enzymes we performed long-time (1.5 μs) equilibrium molecular dynamics for generating conformations starting from holo-conformation of enzyme, *i.e* a ligand bound structure, but ligand is removed before simulation. Each simulation trajectory, subsequently, was clustered based on the distances among closest heavy atoms of BSR. On these clustered structures, we computed the native functionality score to assess the closeness of these to native conformations and deviation from the native state.

## Molecular Dynamics (MD) simulation

We performed explicit water equilibrium Molecular Dynamics simulation on the enzyme structure (using NAMD (Phillips et al., 2005) (Best et al., 2012). Before simulation, missing side chains atoms in tertiary structure were fixed using Foldx(v4) (Schymkowitz et al., 2005). Subsequent to this, the input protein structures were created for simulation using the *autopsfgen* plugin from VMD (Humphrey, Dalke, & Schulten, 1996) and solvated with TIP3P water model using solvate plugin of VMD. The protein box size was extended such that the solvent forms 20 Å thickness around the protein structure. The system was neutralized using the *autoionize* plugin.

The MD simulations were performed using NAMD (v3.0) using CHARMM36 topology and force field parameters for proteins having CMAP correction (Huang & MacKerell, 2013). The protein TIP3P solvated system was minimized for 10,000 steps followed by ∼500 ps temperature equilibration at 300K and pressure equilibration until average pressure of 1 atm was achieved. The temperature was gradually increased in steps of 30 K/2ps to 300K before further equilibration of 500 ps at 300 K. The system was simulated in periodic boundary conditions with electrostatics interactions computed using Particle Mesh Ewald (PME) (Darden, York, & Pedersen, 1993) method by specifying appropriate grid sizes for each solvated protein system. The van der Waals interaction involved switching functions with a cut-off distance of 12 Å (starting at 10 Å). The constant pressure at ∼1 atm was maintained using Nosé-Hoover Langevin piston method (Feller, Zhang, Pastor, & Brooks, 1995) with piston period of 100 fs, a damping timescale of 50 fs and piston temperature of 300 K. A constant temperature of 300 K was maintained using the Langevin dynamics (Paterlini & Ferguson, 1998), with the damping coefficient set to 5 ps^-1^ for all heavy atoms.

A time-step of 2 fs was used in both equilibration and production runs. In the production run of simulation, the trajectory was stored at every 5 ps, which resulted in 3,00,000 frames. The total energies of the simulated systems were stable after the equilibration period and visual inspection of trajectories from the production phase did not show any unfolding for most of the proteins. The trajectories were visualized and analyzed using the VMD (Humphrey et al., 1996) program and Bio3D package (package in R) (Grant, Skjaerven, & Yao, 2021). The plots were prepared using the ggplot2 library (Wickham, 2011) in R.

To generate smoothened representations of structures and computing RMSF, we obtained trajectories after taking stride of 10, which resulted in 30,000 frames. These were superposed on the native structure. We selected loop regions (having high RMSF values) for their relative motion along the trajectory using VMD. For visual smoothened representation, frames were taken at a stride of 100 and smoothened by the value of 25 in VMD.

## Clustering of structures

To analyze a large number of structures for each enzyme, we relied on dimensionality reduction methods to cluster structures based on binding site features. Here, we have defined interatomic distances of heavy atoms of BSR as features for further analyses. Therefore, structures having similar distances among BSR will be conformationally similar and can be clustered together. For this, we relied on methods implemented in the MSM-builder package (Harrigan et al., 2017) package for featurization and clustering (Gao & Zhao, 2017). First, the MD trajectories were converted into vectors containing feature space of interest, referred to as the featurization step, in which we used the minimum distance between heavy atoms of all possible BSR. The number of features varied for each enzyme. These features were used for dimensionality reduction using a method called time-structure independent component analysis (tICA) (Harrigan et al., 2017; Nguyen, 2007).

These independent components represent distinct modes of motion or structural changes including its time component. The tICA could discern independent components or modes capturing the essential slow dynamics present in the input feature space. The tICA method was used with 5 number of components for each enzyme. Subsequently, conformations were clustered using “MiniBatchKMediods” clustering algorithm and visualized on the first two components of tICA with fixed number of clusters. The centroid of the cluster is considered as its representative member.

## Sampling of clusters

To compare the clusters, we randomly drew 100 conformations from each cluster and computed their mean property (RMSD/*fχ*). This process was repeated 100 times for each cluster. This approach covers the general behaviour of the cluster as with the repeated sampling with their mean computation. To assess structural similarity using RMSD, the sampled conformations’ structures (Cα atoms of BSR) were superposed on the first frame (Cα atoms of BSR) of MD simulation. The intra-cluster RMSDs of BSR were calculated after superposing the members of the cluster on their respective centroid or representative.

For generation of the cluster specific loop movement, all the sampled frames (10,000) in the sampling step were superimposed on to the native structure. For the pictorial representation, stride of 10 with a smoothening step of 5 was used to prepare the inset figures of the clusters in panel C of all the figures.

## Native functionality analysis

To assess the binding site similarity of clustered structures to the native structure, we have devised a metric, called **Native functionality (*fχ)* score**, which is computed for each frame. This score is a modified form of the Cai score developed for calculating the functional state of a protein in folding simulation in the work of Nissley et al. (Nissley et al., 2022), The native functionality score is given by equation 1:

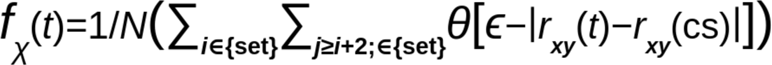

where, the score *fχ* (t) is the fraction of pairwise distances, which are native-like for the given frame (t); the ‘set’ is the list of BSR of an enzyme; indices i and j correspond to residues in ‘set’ of the protein; the N is the total number of pairwise contacts (≤4.5 Å) between heavy atoms of residues where i and j satisfy the condition j≥i+2. The value of ε is taken as 0.2⋅r_xy_, where r is the sum of van der Waals radii of atom x and y (x and y corresponds to the type of heavy atoms of the residues i and j) plus 1Å. The parameters r_xy_(t) and r_xy_(cs) are the distances between atom pair x and y at time t and between same atoms x and y of residues i and j in the native structure, respectively. θ(x) is the step function given by equation (2):

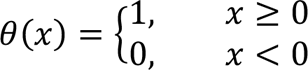

The numerical value of *fχ* ranges from 0 to 1, where the value of 1 is achieved when distances between all heavy atom pairs of the binding site are within tolerated limit (∊) of distances for native structure (‘native structure’). Alternatively, higher value of *fχ* would correspond to conformation similar to native and lower values of *fχ* would be indicative of conformations away from it.

Being a step function, equal weightage is given to all the atomic contacts and value of [0,1] signifies the fraction of contacts that remained within the allowed limits. The way this score is calculated, even a flip of the side chain would lead to atomic distances exceeding the permissible limit thus being reflected in the score. Thus, this score is highly sensitive for relative spatial positioning of atoms. In computation of native functionality of the clusters, we have extracted 100 conformations from a cluster by random sampling for computing *fχ* and subsequently calculating average < *fχ* > of that sample. This process was repeated 100 times to calculate the distribution of <*fχ* > for any given cluster. The dotted line representing the mean of cluster in panel F of the figures represents the mean of this distribution (n=100).

## Acknowledgments

We acknowledge computing facility Param Smriti formed under National Supercomputing Mission (NSM). We acknowledge IISER Mohali and Bioinformatics Center (BT/PR40419/BTIS/137/36/2022) grant from Department of Biotechnology under the Ministry of Science and Technology, Govt. of India for funding and fellowship support.

