## Supporting_information for "Deciphering binding site conformational variability of substrate promiscuous and specialist enzymes"

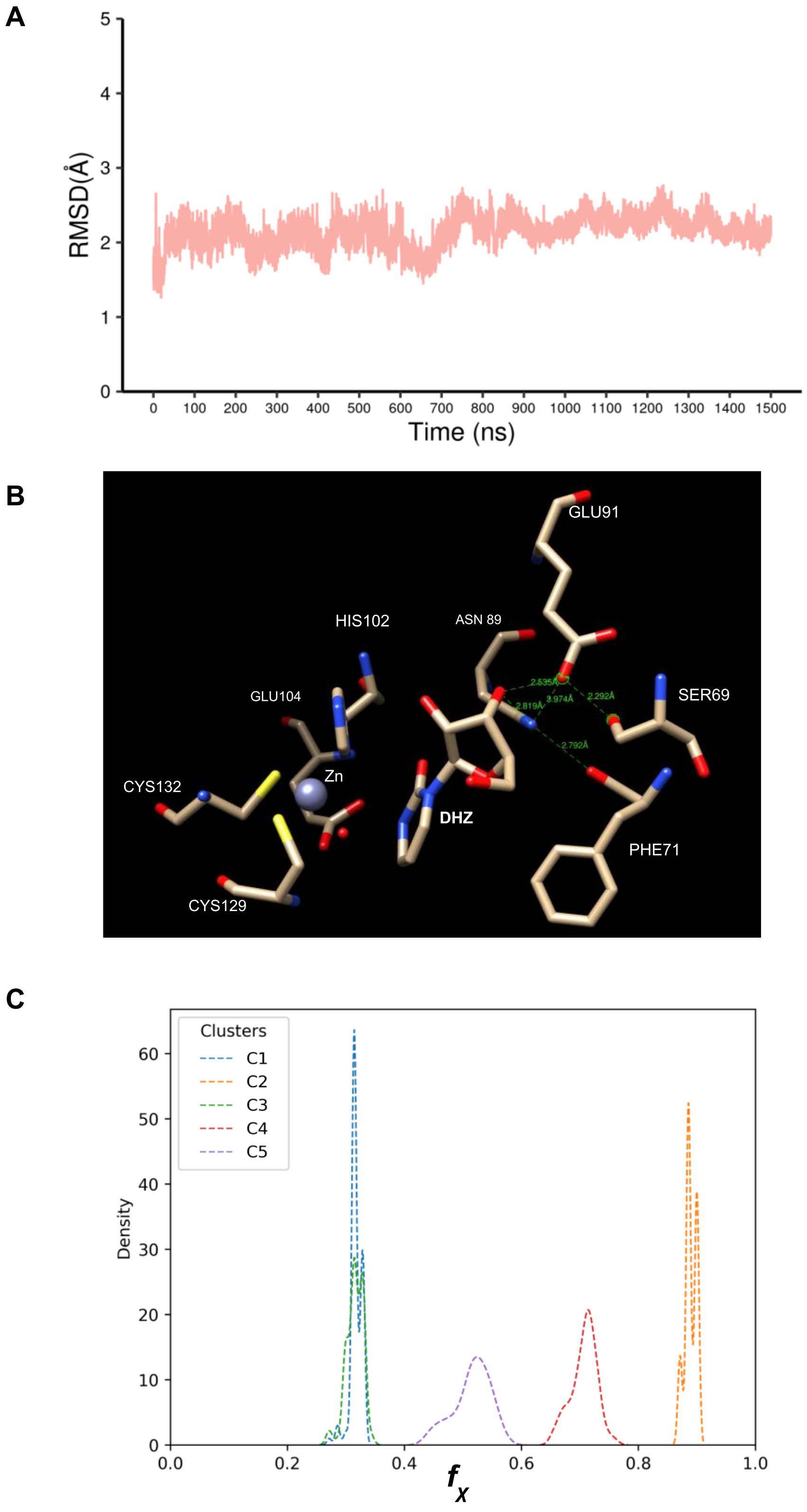


**Figure S1: Analysis of *E. coli* cytidine deaminase enzyme: (A)** Line plot of RMSD versus simulation production time (in ns) (B) Arrangement of a region of binding site highlighting interatomic distances between loop residues S69 and F71 with N89 and E91 residues. (C). Density distribution of mean native functionality ($f_\chi$) of each cluster sampled members

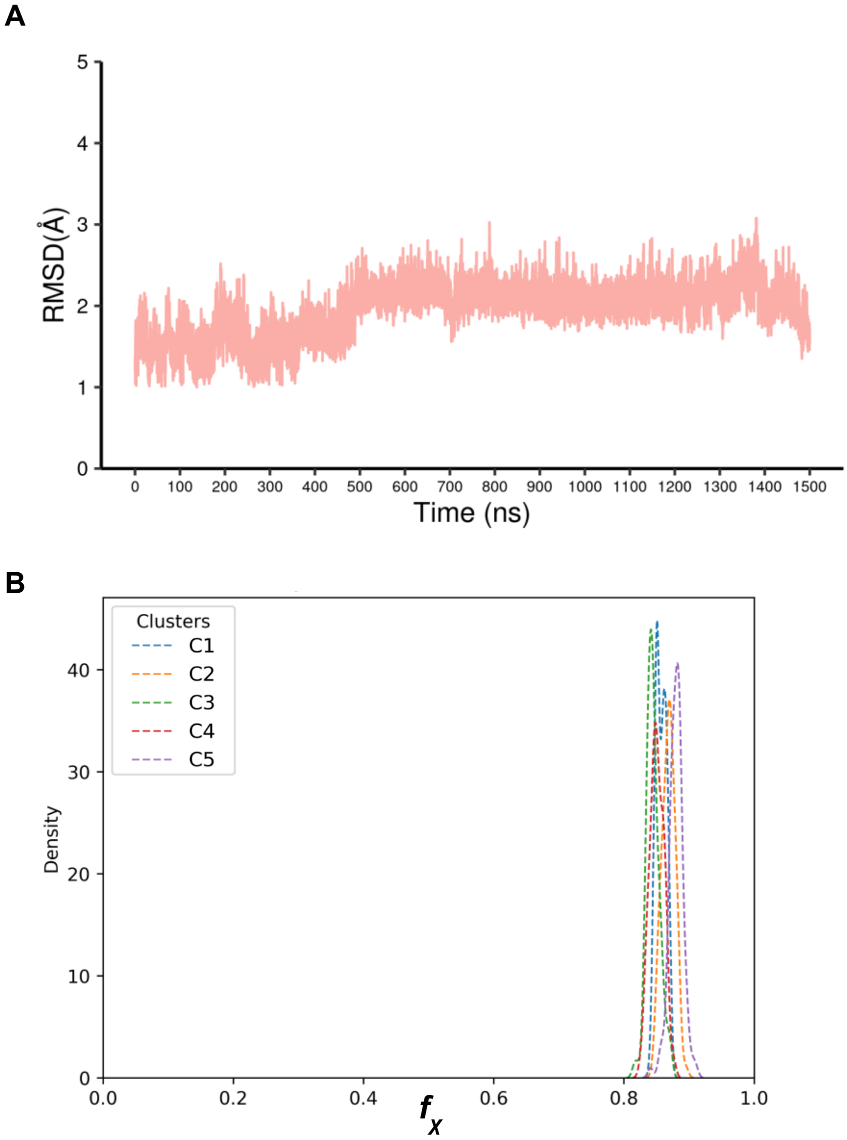


**Figure S2: Analysis of human Dihydrofolate reductase: *(*A)** Line plot of RMSD versus simulation production time (in ns) **(B)**. Density distribution of mean native functionality ($f_\chi$) of each cluster sampled members.

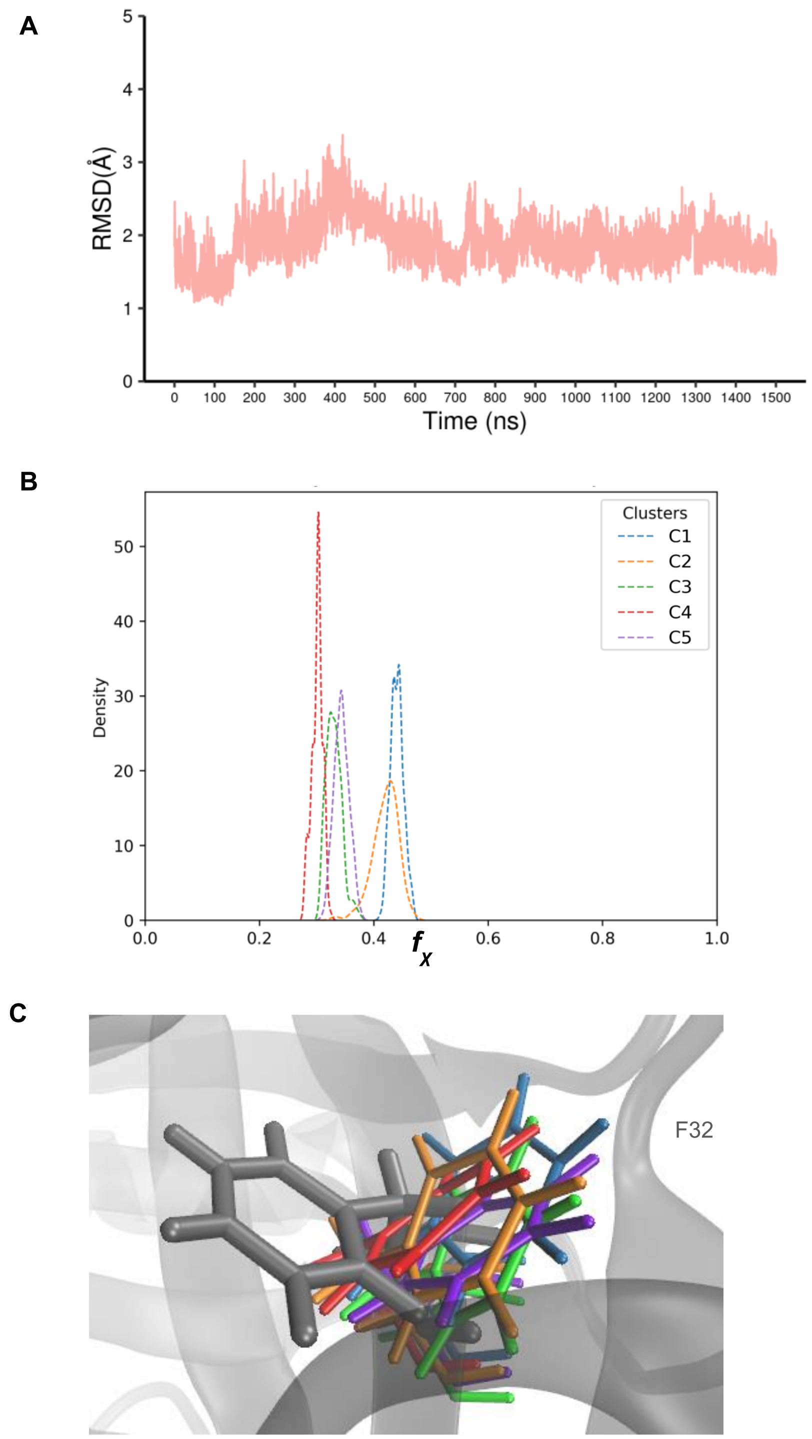


**Figure S3: Analysis of *E. coli* 4-diphosphocytidyl-2-C-methyl-D-erythritol kinase (CDP-ME Kinase): (A)** Line plot of RMSD versus simulation production time (in ns) **(B)** Density distribution of mean native functionality ($f_\chi$) of each cluster sampled members. **(C)** Side chain orientation shown of a binding site residue F32 from all five cluster representatives compared to native structure. Side chain is shown in licorice representation with native shown in grey color, while rest others cluster residues are shown in different color.

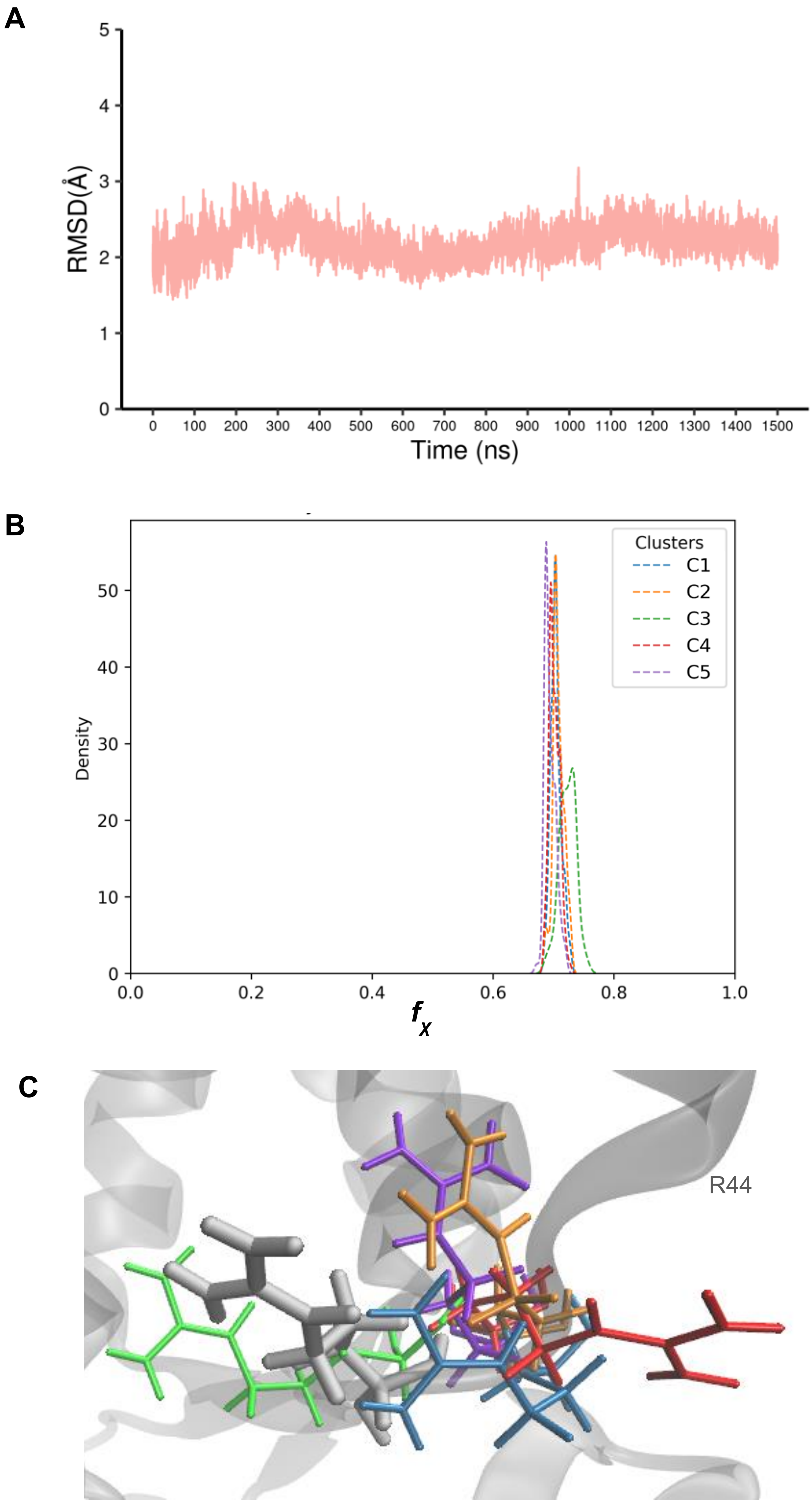


**Figure S4: Analysis of *Psuedomonas aeruginosa* p-hydroxybenzoate hydroxylase (*PBH*):** Line plot of RMSD versus simulation production time (in ns) **(B**) Density distribution of mean native functionality ($f_\chi$) of each cluster sampled members. **(C)** Side chain orientation shown for binding site residue R44 of native and cluster representative structures. Side chain is shown in licorice representation and native is colored grey while rest others are different colored.
